# With or without a Ca^2+^ signal? A proteomics approach towards Ca^2+^ dependent and independent proteome changes in response to oxidative stress in *A. thaliana*

**DOI:** 10.1101/2025.03.31.645912

**Authors:** Annelotte van Dieren, Andras Bittner, Bernhard Wurzinger, Leila Afjehi-Sadat, Wolfram Weckwerth, Markus Teige, Ute C. Vothknecht

## Abstract

Calcium (Ca^2+^) and reactive oxygen species (ROS) are key secondary messengers in plant stress signaling, yet their interplay in regulating proteome-wide responses remains poorly understood. In this study, we employed label-free quantitative (LFQ) proteomics to investigate Ca^2+^-dependent and independent changes in the proteome of *Arabidopsis thaliana* leaves upon oxidative stress induced by hydrogen peroxide (H_2_O_2_). To dissect the role of Ca^2+^ signaling, we inhibited H_2_O_2_-induced Ca^2+^ transients by pretreatment with LaCl_3_, a plasma membrane Ca^2+^ channel blocker. We then analysed the proteome of plants treated with H_2_0_2_ or ddH_2_O after 10 and 30 min of treatment and detected 3724 and 3757 proteins, respectively. From these, 581 proteins showed significant changes in abundance after 10 min and 909 proteins after 30 min. Remarkably, the combined LaCl_3_ and H_2_O_2_ treatment resulted in the highest number of differentially abundant proteins (DAPs), indicating a strong attenuating effect of Ca^2+^ signaling on the oxidative stress response. Specifically responsive to only H_2_O_2_ were 37 and 57 proteins with distinct subsets of strictly Ca^2+^-dependent, partially Ca^2+^- dependent, and Ca^2+^-independent proteins. Notably, Ca^2+^-independent H_2_O_2_-responsive proteins predominantly showed increased abundance, while strictly Ca^2+^-dependent proteins exhibited decreased abundance, suggesting a role for Ca^2+^ signaling in protein degradation. Furthermore, three proteins—WLIM1, CYP97C1, and AGAP1—underwent Ca^2+^-dependent shifts between the two time points, pointing to a dynamic nature of Ca^2+^-regulated proteomic changes. This study provides novel insights into short-term Ca^2+^-dependent and independent regulation of the Arabidopsis leaf proteome in response to oxidative stress, identifying key stress-responsive proteins and potential new targets for further research on plant stress resilience mechanisms.

## Introduction

Plants are continuously exposed to various environmental stresses, including drought, salinity, extreme temperatures, and pathogen attacks. These stressors can disrupt cellular homeostasis, leading to the overproduction of reactive oxygen species (ROS), which can cause oxidative damage to cellular components such as lipids, proteins, and nucleic acids (Mittler, 2017). However, plants have evolved sophisticated signaling networks to perceive and respond to oxidative stress, with Ca^2+^ signaling playing a central role in orchestrating these adaptive responses (Li et al., 2022). Calcium ions (Ca^2+^) serve as a ubiquitous second messenger in plant cells, regulating various physiological and developmental processes. A diverse array of biotic and abiotic stress factors, along with various developmental processes, can induce increases in cytosolic calcium concentration [Ca^2+^]_cyt_ through a regulated influx of Ca^2+^ from both extracellular sources and intracellular reservoirs into the cytosol (McAinsh & Pittman, 2009; Kudla et al., 2010). These transient elevations in [Ca^2+^]_cyt_ exhibit distinct spatio-temporal characteristics, including variations in amplitude, frequency, and subcellular localization, in a manner that is specific to the type of stimulus encountered. The unique patterns of [Ca^2+]^_cyt_ fluctuations, commonly referred to as “calcium signatures,” (Allen et al., 2001; Whalley & Knight, 2013) play a crucial role in ensuring the specificity of calcium-mediated signaling, thereby facilitating context-dependent and stimulus-appropriate cellular responses. Each calcium signature arises from the coordinated and dynamic interplay of multiple calcium influx channels and efflux transporters, which are located within the plasma membrane as well as the membranes of various intracellular organelles (Demidchik et al., 2018). These external stimuli are decoded by calcium-binding proteins, including calmodulins (CaMs), calcium-dependent protein kinases (CDPKs), and calcineurin B-like proteins (CBLs), which send the signal to downstream effectors (Mohanta et al., 2019; Tang et al., 2020). Furthermore, different plant organs and tissues exhibit distinct calcium signatures in response to stress, emphasizing the complexity and specificity of Ca^2+^-mediated signaling networks (Costa et al., 2018; Giridhar et al., 2022).

After the initial identification of stimulus-specific changes in [Ca^2+^]_cyt_, an increasing number of processes involving Ca^2+^ signaling have been elucidated, including those related to plant growth and development, such as cell division and organ formation (Zhang et al., 2014). Numerous studies have investigated a variety of calcium inducing stimuli across different plant species, including NaCl, mannitol, H_2_O_2_, and Flg22 in barley leaf and root samples (Giridhar et al., 2022), and NaCl, mannitol, H_2_O_2_ and Pep13 in potato (Van Dieren et al., 2024). Additionally, downstream responses controlled by these specific Ca^2+^ signals have been described, including the role of Ca^2+^-regulated kinases in mediating phosphorylation events that coordinate signaling cascades (Ludwig, 2003) as well as responses that comprise regulation of gene expression through Ca^2+^-regulated transcriptional responses (Kaplan et al., 2006) and Ca^2+^-responsive promotor elements (Kudla et al., 2010). Oxidative stress results from an imbalance between ROS production and detoxification. While excessive ROS can be detrimental, controlled ROS production acts as a signaling molecule that activates stress-responsive pathways (Mittler, 2017; Chen & Yang, 2020). Rapid signaling and communication from individual cells that perceive potential threats to their neighbouring cells as well as more distal tissue is vital for plant acclimation and fitness. In this context, it was shown that calcium signaling and ROS interact in a complex feedback loop. ROS can induce Ca^2+^ influx through plasma membrane and organellar channels, leading to further signal propagation (Li et al., 2022; Ravi et al., 2023). In turn, Ca^2+^ signaling modulates ROS-scavenging mechanisms, such as the activation of antioxidant enzymes including superoxide dismutase (SOD), catalase (CAT), and ascorbate peroxidase (APX) (Gilroy et al., 2016). Other studies highlight the role of NADPH oxidases, also known as respiratory burst oxidase homologs (RBOHs), in ROS production upon Ca^2+^ signaling activation (Kärkönen & Kuchitsu, 2015). These enzymes facilitate ROS bursts that act as secondary messengers, amplifying stress responses. Moreover, Ca^2+^ channels such as cyclic nucleotide-gated channels (CNGCs) and glutamate receptor-like channels (GLRs) contribute to ROS-Ca^2+^ crosstalk, further fine-tuning the stress response (Gilroy et al., 2016). The interplay between calcium signaling and oxidative stress represents a crucial aspect of plant stress responses. Understanding these mechanisms provides insights into how plants adapt to adverse environmental conditions and could be exploited for the development of stress-resilient crops.

One of the first layers of cellular signaling is the translation of secondary signal components into re- adjustments of the transcriptional machinery. Consequently, many large-scale approaches to study stress responses analyse changes in gene expression. However, proteins are key players in the structure, function, and regulation of cells, tissues, and organs, and proteome changes can occur independent from transcription by processes such as protein degradation and regulation of translation (Gry et al., 2009; Payne, 2015; Liu et al., 2016). The interplay of ROS and Ca^2+^ signaling on transcriptome changes have recently been investigated in barley (Bhattacharyya et al., 2025), however, no investigation has so far described the effect of Ca^2+^ signaling on ROS induced changes of proteomes. We thus aimed to elucidate the role of H_2_O_2_-induced Ca^2+^ signals on short-term proteome changes observed in Arabidopsis leaf tissue by inhibiting stress induced Ca^2+^ transients using the established plasma membrane Ca^2+^ channel blocker LaCl3 (Tracy et al., 2008). MS-based proteome analysis identified specific subsets of proteins, whose abundance changed upon 10 and 30 min H_2_O_2_ application in a Ca^2+^-dependent or -independent manner. However, one of the major challenges in omics is translating high-dimensional data into meaningful biological insights with practical applications. The integration of proteomic data with other omics datasets (e.g., genomics, transcriptomics, and understanding of complex biological processes. This knowledge could be further investigated and applied to future research aiming to enhance stress resistance and optimizing performance and productivity in crop species under increasingly challenging environmental conditions.

## Results

H_2_O_2_ and Ca^2+^ are secondary messengers that are involved in the mediation of environmental changes into an appropriate cellular response. Temporal increases in these messengers affect various cellular processes including gene transcription or protein activity. Here we employed label free quantitative (LFQ) proteomics to analyse the H_2_O_2_ induced changes in the leaf proteome of Arabidopsis and the contribution of Ca^2+^ signals in the H2O_2_ induced changes.

### Establishing parameters and experimental design

H_2_O_2_ induced Ca^2+^ transients and their inhibition by the Ca^2+^ channel blocker La^3+^ have been shown before for Arabidopsis (Giridhar et al., 2022; Rentel & Knight, 2004; Van Dieren et al., 2024). To confirm that these responses also occur under the experimental conditions chosen for the protein isolation, leaf discs from 3-week-old At-AEQ_cyt_ plants grown under the same circumstances as wild type plants used for proteomics analysis were analysed. As shown before for soil-grown Arabidopsis plants (Van Dieren et al., 2024), an oxidative stress stimulus of 20 mM H_2_O_2_ resulted in a well-shaped Ca^2+^ transient in Arabidopsis leaf tissue, which was inhibited by over 50 % upon pre-treatment with 1 mM LaCl_3_ (Fig. 1A).

**Fig. 1.**
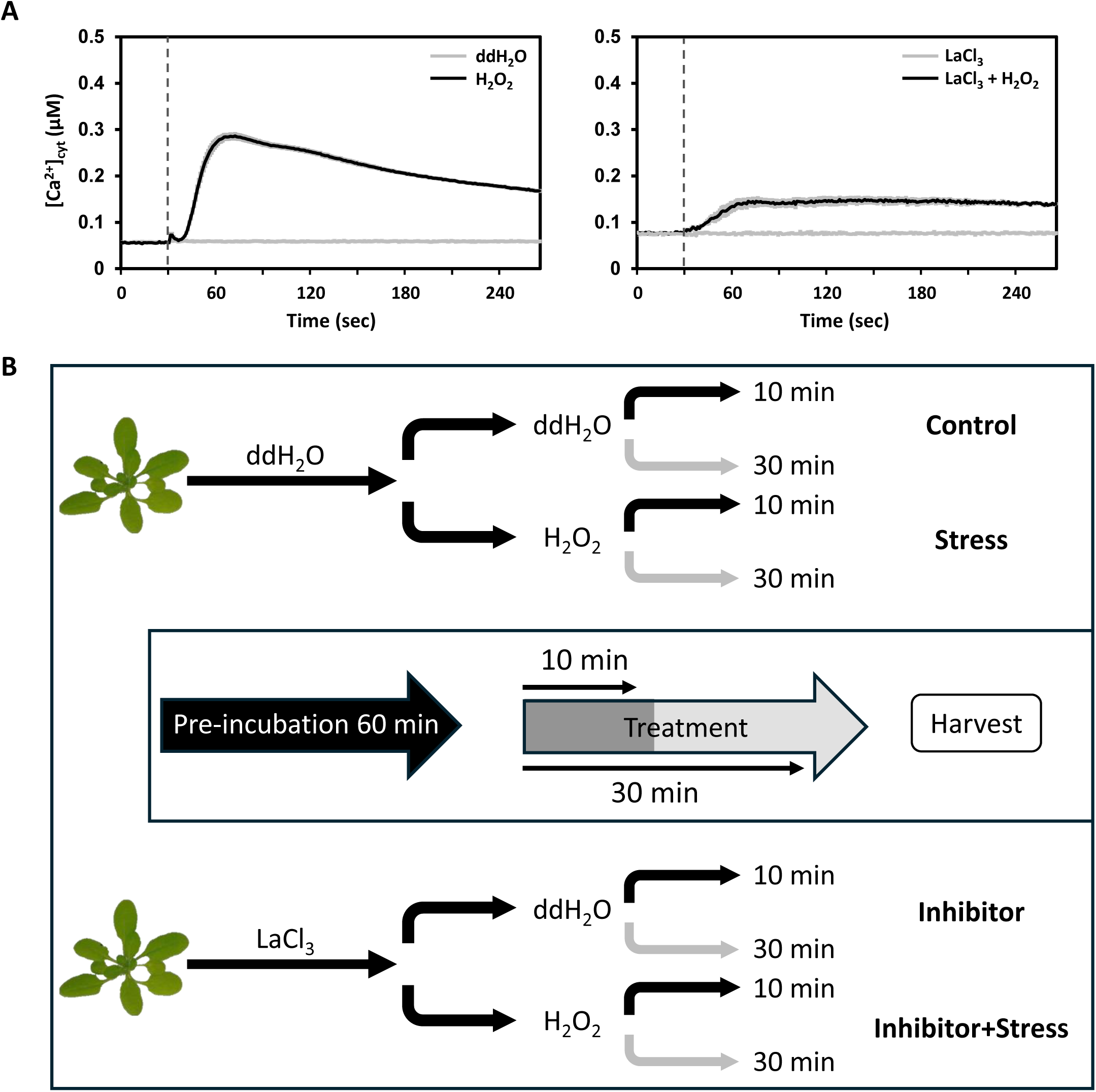
Experimental background and design. (**A**) Time course of changes in [Ca^2+^]_cyt_ in response to 20 mM H_2_O_2_ in leaf tissue of Arabidopsis (left) and in response to 20 mM H_2_O_2_ after 60 min pre-incubation with LaCl_3_ (right). Values are shown as mean ± SE (n = 6). Dashed vertical lines indicate the time point of stimuli injection (30 sec). (**B**) Overview of treatment application: plants were either pre-incubated in ddH_2_O or 1 mM LaCl_3_ (Inhibitor) for 60 min. Half of the plants from both pre-incubations were transferred into a 20 mM H_2_O_2_ solution (Stress), another half was transferred into fresh ddH_2_O. Half of these plants were harvested after 10 minutes, the other half after 30 minutes and labelled as indicated: Control (ddH_2_O + ddH_2_O), Stress (ddH_2_O + 20 mM H_2_O_2_), Inhibitor (1 mM LaCl_3_ + ddH_2_O), Inhibitor + Stress (1 mM LaCl_3_ + 20 mM H_2_O_2_) with their corresponding treatment duration (10 or 30 min).

The workflow of the application of the different treatments before proteomics analyses is schematically displayed in Fig. 1B. Complete rosettes from 3-week-old wild type plants grown on soil were pre-incubated with LaCl3 (Inhibitor) or ddH_2_O for 60 min. Subsequently, the rosettes were washed carefully and treated with either 20 mM H_2_O_2_ (Stress) or ddH_2_O for 10 and 30 min. The timing was chosen to elucidate short-term responses to the stress stimulus. The different treatment paths result in the following treatment names used further: Control: pre-incubation in ddH_2_O, treatment with ddH_2_O; Stress: pre-incubation with ddH_2_O, treatment with H_2_O_2_; Inhibitor: pre-incubation with LaCl_3_, treatment with ddH_2_O; Inhibitor+Stress: pre-incubation with LaCl_3_, treatment with H_2_O_2_.

### Initial data analysis

For each of the four different treatments (Control, Stress, Inhibitor and Inhibitor+Stress), five independent biological replicates, each consisting of pooled proteins from 12 rosettes, were analysed.

The proteome analysis of the samples resulted in the identification of 3724 proteins after 10 min and 3757 proteins after 30 min of stress treatment (Fig. 2A). The number of identified proteins is in line with former proteomics analyses in Arabidopsis (Seaton et al., 2018; Ayash et al., 2021; Scholz et al., 2025). For the further analysis we separated the LFQ intensities in two groups: one group representing the samples harvested after 10 min of stress treatment, the other group representing the samples harvested after 30 min of stress treatment. After a quality control step, in which proteins ‘only identified by site, reverse sequences, and potential contaminants’ were filtered out, 2906 proteins remained for the 10 min samples and 2965 proteins for the 30 min samples (Fig 2A, quality control). After multiple sample ANOVA test (p-value <0.05), 581 proteins remained for the 10 min samples and 909 for 30 min (Fig. 2A, statistical analysis). To ensure biological validity, stringent filtering is applied, leading to significant dataset reduction.

**Fig. 2:**
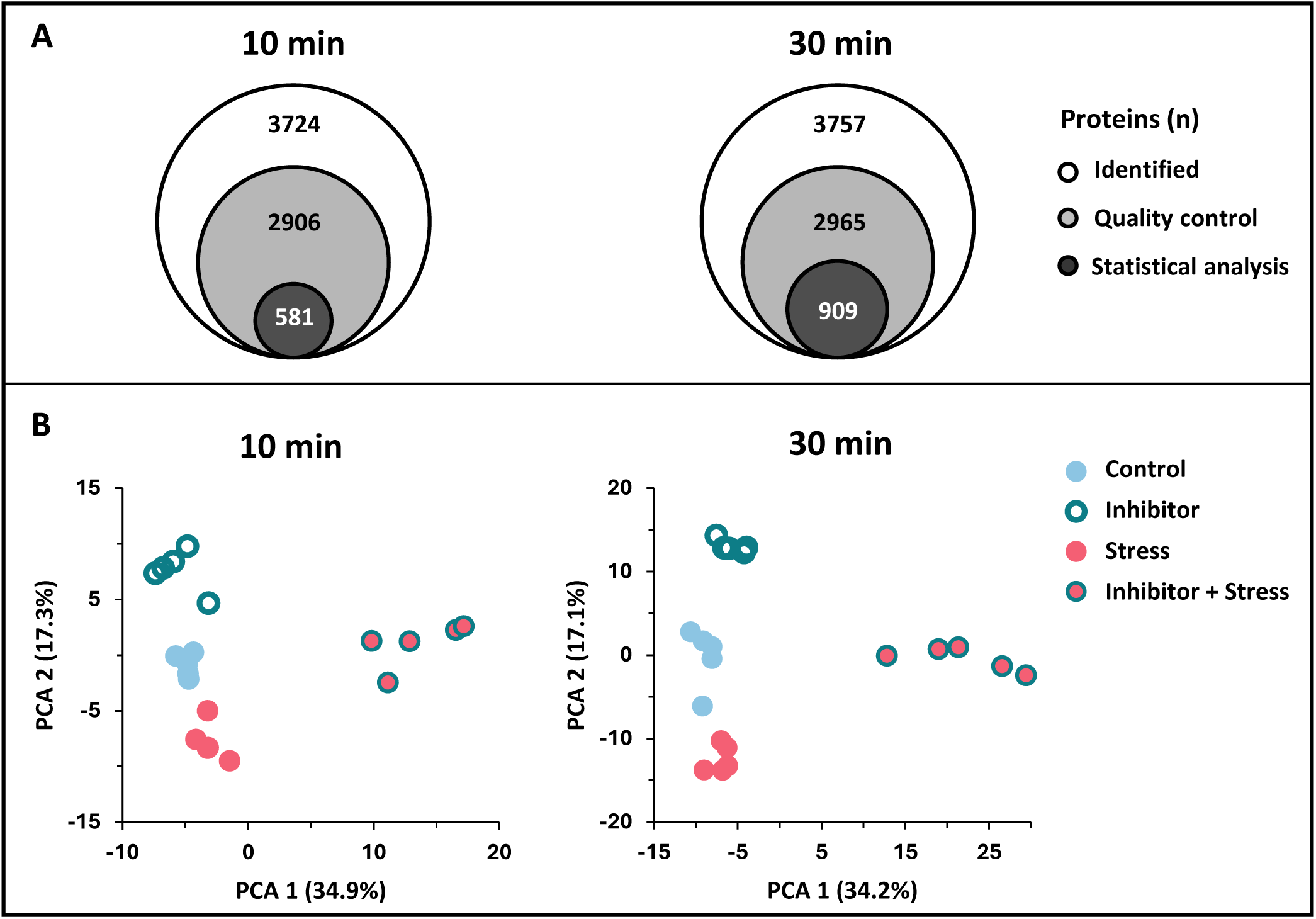
Overview of the initial data analysis for the 10 min (left) and 30 min (right) samples with (**A**) showing the numbers of all proteins identified (identified), proteins left after removal of contaminants, proteins only identified by site, and reverse annotated peptides (quality control), and proteins left after statistical analysis (ANOVA, p≤0.05). (**B**) Principal component analysis (PCA) of the LFQ intensities of the quantified proteins, colours indicate the different treatments: blue fill = Control, magenta cycle = Stress, turquoise fill = Inhibitor and magenta cycle + turquoise fill = Inhibitor+Stress.

A principal component analysis (PCA) of the LFQ values showed that all replicates clearly fell into their corresponding treatment group (Fig. 2B). PC1 explains more than 34% of the variance within the data for both time points and essentially separating the Inhibitor+Stress treatment from all other treatments, while PC2 (>17 % of the variance for both time points) separates the Control, Stress and Inhibitor treatments. Overall, this analysis indicates clear proteomic changes, especially with regards to the Inhibitor+Stress samples compared to all other treatments. The other treatments cluster closer together but still remain separated from each other.

### Clustering analysis

In line with the PCA analysis, the hierarchical clustering of protein abundance, performed on the Z-scored normalized intensities, showed a clear clustering of all replicates of an individual treatment (Fig. 3A, dendrograms on top of the heatmaps) and therewith the strong similarity in protein abundance among the replicates within one treatment group. All five replicates of the Inhibitor+Stress treatment clustered together as one of two main clusters. In the other main cluster, two subclusters could be observed, with the first one representing the 5 replicates of the Stress treatment, and the second one showing a close relation of the Control and Inhibitor only treatment. This pattern of clustering was observed for both the 10 and the 30 min of stress treatment. It substantiates the strong effect of the Inhibitor+Stress treatment on the proteome, and the slightly milder but clear effect of the Stress treatment alone, while the Inhibitor treatment alone has a lesser effect. It also indicates an attenuating effect of Ca^2+^ signaling on the H_2_O_2_-induced stress response.

**Fig. 3:**
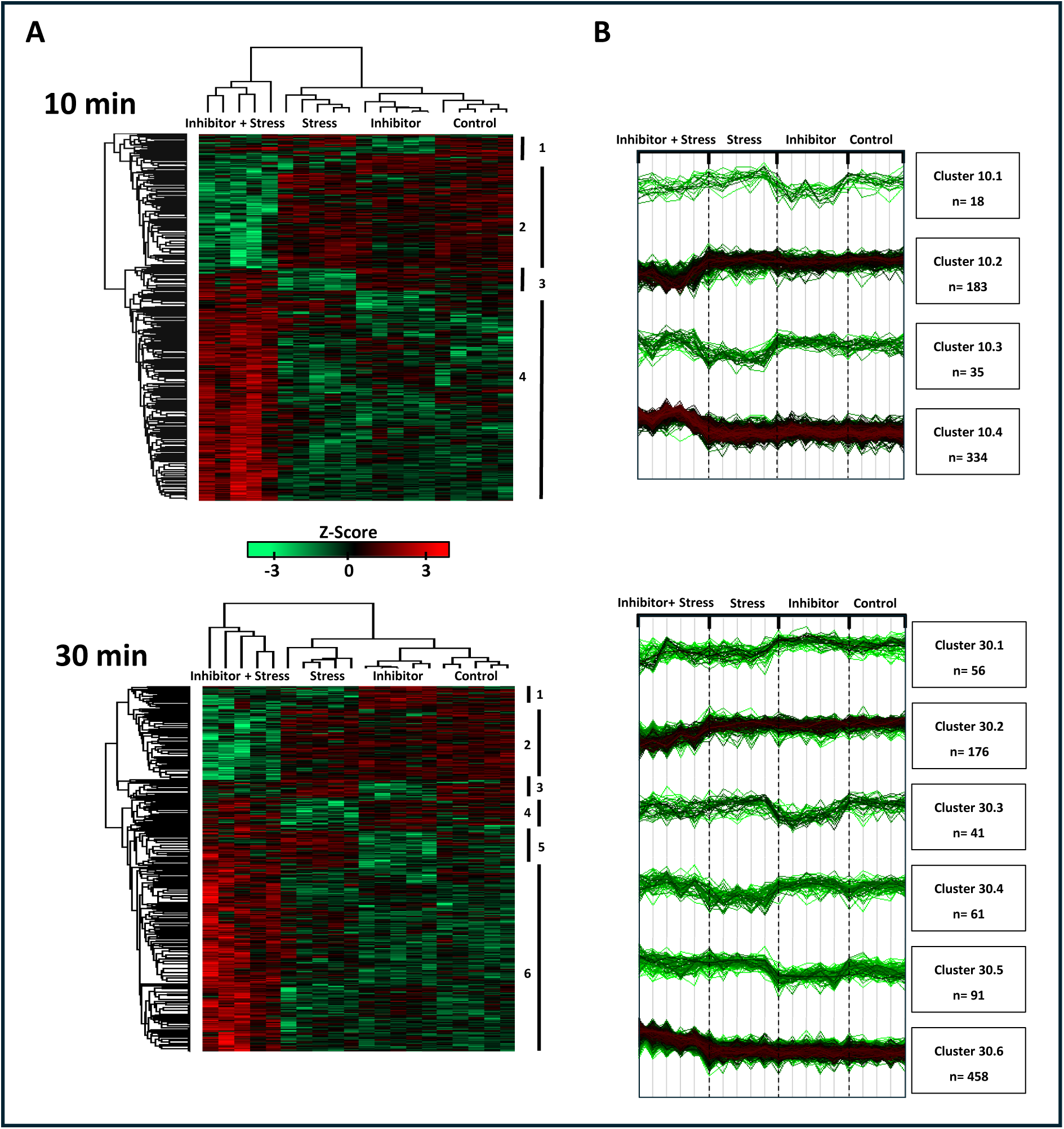
Heatmaps and hierarchical clustering of treatments and identified DAPs. **A**: Heatmaps showing the z-score overall pattern of relative increased (red) and decreased (green) protein abundance within the samples after 10 (upper panel) and 30 (lower panel) min of treatment. The dendrogram of the columns (top) shows how the four different treatments separate based on Euclidean distance. The dendrogram of the rows (left side) shows the clustering of the protein. Identified protein clusters are indicated with a number on the right side of both heatmaps. **B**: Intensity plots for each cluster from panel A and number of proteins (n) in the specific cluster is indicated.

Hierarchical clustering further revealed segregation of the proteins into 4 different abundance clusters after 10 min of treatment, and 6 clusters after 30 min of treatments (Fig. 3A, dendrograms on the side of the heatmaps). The differences in protein abundance of the different samples are shown in the heatmap with green colour indicating a decrease and red colour an increase in relative protein abundance (Fig. 3A) as well as profile plots (Fig. 3B). The 10 min clusters were characterized by a lower protein abundance for the Inhibitor and Inhibitor+Stress treatment (cluster 10.1), a lower abundance for the Inhibitor+Stress treatment (cluster 10.2), a lower abundance for the Stress treatment (cluster 10.3) and a lower abundance for all but the Inhibitor+Stress treatment (cluster 10.4). The clusters identified after 30 min of treatment were defined by a lower abundance for the Inhibitor+Stress and the Stress treatment (30.1), a lower abundance for the Inhibitor+Stress treatment (30.2), a lower abundance for the Inhibitor treatment (30.3), a lower abundance for the Stress treatment (30.4), a lower abundance for the Control and Inhibitor treatment (30.5), and a lower abundance for all but the Inhibitor+Stress treatment (30.6). At both time points the clusters containing proteins with different abundance in the Inhibitor+Stress treatment only were the largest clusters, containing 183 and 334 proteins for the 10 min (10.2 and 10.4), and 176 and 458 proteins for the 30 min (30.2 and 30.6).

### Gene Ontology analysis of heatmap clusters

Proteins in each cluster were functionally classified by Gene Ontology (GO) and KEGG term enrichment analysis. Details on the different GO-terms (molecular function and biological process) of all clusters are shown in Table 1 (10 min) and Table 2 (30 min). Enriched terms in three out of four clusters with a relative lower abundance of proteins in the Inhibitor+Stress treatment (10.1, 10.2 and 30.2) included Ribosomal pathways. Cluster 10.2, with lower abundance only for Inhibitor+Stress, also included the KEGG term proteasome. Clusters 10.4, 30.5 and 30.6, all of which comprise a relative higher abundance of proteins in the Inhibitor-Stress treatment, have carbon fixation and other pathways related to carbon metabolisms as the most enriched KEGG terms.

**Table 1:** Functional classification of the protein clusters after 10 min of treatment, obtained after hierarchical clustering. For each cluster the top 3 GO-terms (KEGG, molecular function, biological process), selected by FDR and sorted by fold enrichment, are displayed.

| Cluster | Proteins (n) | Analysis | Enrichment FDR | Protein (n) | Pathway protein (n) | Fold Enrichment | Pathways |
| --- | --- | --- | --- | --- | --- | --- | --- |
| 1 | 18 | KEGG | 7.40E-09 | 7 | 315 | 34.1 | Ribosome |
|  |  | GO Molecular function | 3.60E-07 | 7 | 423 | 25.4 | Structural constituent of ribosome |
|  |  |  | 1.00E-06 | 7 | 545 | 19.7 | Structural molecule activity |
|  |  |  | 5.00E-05 | 7 | 1030 | 10.4 | MRNA binding |
|  |  | GO Biological Process | 5.40E-03 | 6 | 934 | 9.9 | Amide biosynthetic proc. |
|  |  |  | 6.30E-03 | 6 | 1086 | 8.5 | Cellular amide metabolic proc. |
|  |  |  | 8.00E-03 | 7 | 1808 | 5.9 | Organonitrogen compound biosynthetic proc. |
| 2 | 183 | KEGG | 8.90E-38 | 40 | 315 | 19.2 | Ribosome |
|  |  |  | 4.90E-04 | 5 | 61 | 12.4 | Proteasome |
|  |  |  | 3.40E-03 | 4 | 53 | 11.4 | Phenylalanine, tyrosine and tryptophan biosynthesis |
|  |  | GO Molecular function | 1.20E-05 | 3 | 3 | 151 | MAP-kinase scaffold activity |
|  |  |  | 1.20E-05 | 3 | 3 | 151 | Protein kinase C binding |
|  |  |  | 1.20E-05 | 3 | 3 | 151 | Signaling adaptor activity |
|  |  | GO Biological Process | 7.20E-35 | 54 | 860 | 9.5 | Translation |
|  |  |  | 7.20E-35 | 54 | 865 | 9.4 | Peptide biosynthetic proc. |
|  |  |  | 1.90E-34 | 55 | 934 | 8.9 | Amide biosynthetic proc. |
| 3 | 35 | KEGG | 1.0E-03 | 2 | 13 | 121.4 | Sulfur relay system |
|  |  |  | 2.9E-03 | 2 | 25 | 63.1 | Biosynthesis of unsaturated fatty acids |
|  |  |  | 5.7E-03 | 2 | 43 | 36.7 | Alpha-Linolenic acid metabolism |
|  |  | GO Molecular function | 8.7E-04 | 2 | 5 | 315.7 | Acetyl-CoA C-acyltransferase activity |
|  |  |  | 8.7E-04 | 2 | 5 | 315.7 | Thiosulfate sulfurtransferase activity |
|  |  |  | 3.6E-03 | 2 | 15 | 105.2 | Sulfurtransferase activity |
|  |  | GO Biological Process | 1.0E-05 | 4 | 36 | 87.7 | Cysteine metabolic proc. |
|  |  |  | 4.5E-06 | 5 | 75 | 52.6 | Sulfur amino acid metabolic proc. |
|  |  |  | 5.6E-05 | 7 | 437 | 12.6 | Sulfur compound metabolic proc. |
| 4 | 334 | KEGG | 8.00E-26 | 23 | 69 | 27.6 | Carbon fixation in photosynthetic organisms |
|  |  |  | 1.50E-14 | 16 | 77 | 17.2 | Glyoxylate and dicarboxylate metabolism |
|  |  |  | 4.30E-11 | 12 | 58 | 17.1 | Pentose phosphate pathway |
|  |  | GO Molecular function | 4.10E-08 | 6 | 10 | 49.6 | L-malate dehydrogenase activity |
|  |  |  | 1.40E-07 | 7 | 20 | 29 | Intramolecular oxidoreductase activity, interconverting aldoses and ketoses |
|  |  |  | 1.10E-09 | 18 | 180 | 8.3 | Oxidoreductase activity, acting on CH-OH group of donors |
|  |  | GO Biological Process | 7.60E-31 | 44 | 317 | 11.5 | Response to cadmium ion |
|  |  |  | 4.70E-18 | 32 | 315 | 8.4 | Nucleotide metabolic proc. |
|  |  |  | 4.30E-36 | 78 | 1055 | 6.1 | Carboxylic acid metabolic proc. |

**Table 2:** Functional classification of the protein clusters after 30 min of treatment, obtained after hierarchical clustering. For each cluster the top 3 GO-terms (KEGG, molecular function, biological process), selected by FDR and sorted by fold enrichment, are displayed.

| Cluster | Proteins (n) | Analysis | Enrichment FDR | Protein (n) | Pathway protein (n) | Fold Enrichment | Pathways |
| --- | --- | --- | --- | --- | --- | --- | --- |
| 1 | 56 | KEGG | 1.90E-03 | 3 | 61 | 24.3 | Citrate cycle (TCA cycle) |
|  |  |  | 1.10E-03 | 4 | 119 | 16.6 | Glycolysis / Gluconeogenesis |
|  |  |  | 1.10E-03 | 4 | 120 | 16.4 | Cysteine and methionine metabolism |
|  |  | GO Molecular function | 1.90E-04 | 8 | 423 | 9.3 | Structural constituent of ribosome |
|  |  |  | 1.90E-04 | 9 | 545 | 8.1 | Structural molecule activity |
|  |  |  | 4.70E-04 | 9 | 674 | 6.6 | Zinc ion binding |
|  |  | GO Biological Process | 7.90E-04 | 4 | 67 | 29.5 | Aspartate family amino acid metabolic proc. |
|  |  |  | 2.50E-03 | 6 | 317 | 9.3 | Response to cadmium ion |
|  |  |  | 2.50E-03 | 7 | 468 | 7.4 | Cellular amino acid metabolic proc. |
| 2 | 176 | KEGG | 1.30E-35 | 38 | 315 | 18.9 | Ribosome |
|  |  |  | 1.40E-03 | 4 | 41 | 15.3 | Propanoate metabolism |
|  |  |  | 9.80E-07 | 9 | 120 | 11.8 | Cysteine and methionine metabolism |
|  |  | GO Molecular function | 1.90E-19 | 20 | 144 | 21.8 | RRNA binding |
|  |  |  | 3.80E-41 | 46 | 423 | 17.1 | Structural constituent of ribosome |
|  |  |  | 3.80E-41 | 50 | 545 | 14.4 | Structural molecule activity |
|  |  | GO Biological Process | 6.00E-36 | 54 | 860 | 9.9 | Translation |
|  |  |  | 6.10E-36 | 54 | 865 | 9.8 | Peptide biosynthetic proc. |
|  |  |  | 2.30E-37 | 57 | 934 | 9.6 | Amide biosynthetic proc. |
| 3 | 41 | KEGG | 4.30E-02 | 1 | 3 | 224.6 | Caffeine metabolism |
|  |  |  | 4.30E-02 | 2 | 77 | 17.5 | Phagosome |
|  |  |  | 3.60E-02 | 3 | 157 | 12.9 | Endocytosis |
|  |  | GO Molecular function | 1.30E-03 | 5 | 211 | 16 | GTPase activity |
|  |  |  | 3.70E-03 | 5 | 303 | 11.1 | GTP binding |
|  |  |  | 3.70E-03 | 5 | 323 | 10.4 | Guanyl nucleotide binding |
|  |  | GO Biological Process | 3.90E-02 | 2 | 16 | 84.2 | Mitochondrial fission |
|  |  |  | 3.90E-02 | 2 | 18 | 74.9 | Purine-containing compound catabolic proc. |
|  |  |  | 3.90E-02 | 3 | 97 | 20.8 | Endocytosis |
| 4 | 61 | KEGG | 3.30E-03 | 3 | 57 | 23.8 | Aminoacyl-tRNA biosynthesis |
|  |  |  | 9.30E-05 | 5 | 98 | 23.1 | Biosynthesis of nucleotide sugars |
|  |  |  | 1.90E-04 | 5 | 131 | 17.3 | Amino sugar and nucleotide sugar metabolism |
|  |  | GO Molecular function | 2.90E-02 | 3 | 72 | 18.9 | Aminoacyl-tRNA ligase activity |
|  |  |  | 2.90E-02 | 3 | 72 | 18.9 | Ligase activity, forming carbon-oxygen bonds |
|  |  |  | 2.90E-02 | 3 | 76 | 17.9 | Actin filament binding |
|  |  | GO Biological Process | 2.10E-03 | 3 | 19 | 71.5 | Pentose metabolic proc. |
|  |  |  | 7.60E-03 | 5 | 188 | 12 | Monosaccharide metabolic proc. |
|  |  |  | 2.80E-04 | 9 | 468 | 8.7 | Cellular amino acid metabolic proc. |
| 5 | 91 | KEGG | 5.90E-03 | 6 | 269 | 6.8 | Carbon metabolism |
|  |  |  | 2.40E-03 | 14 | 1243 | 3.4 | Biosynthesis of secondary metabolites |
|  |  | GO Molecular function | 8.70E-03 | 3 | 29 | 31.4 | Poly(U) RNA binding |
|  |  |  | 8.70E-03 | 3 | 33 | 27.6 | Poly-pyrimidine tract binding |
|  |  |  | 8.70E-03 | 4 | 82 | 14.8 | NAD binding |
|  |  | GO Biological Process | 1.00E-03 | 6 | 144 | 12.7 | Photosynthesis, light reaction |
|  |  |  | 6.30E-03 | 7 | 317 | 6.7 | Response to cadmium ion |
|  |  |  | 3.90E-04 | 11 | 517 | 6.5 | Generation of precursor metabolites and energy |
| 6 | 458 | KEGG | 2.70E-21 | 22 | 69 | 19.3 | Carbon fixation in photosynthetic organisms |
|  |  |  | 1.90E-17 | 20 | 77 | 15.7 | Glyoxylate and dicarboxylate metabolism |
|  |  |  | 5.70E-13 | 15 | 61 | 14.9 | Citrate cycle (TCA cycle) |
|  |  | GO Molecular function | 7.70E-09 | 8 | 17 | 28.4 | Malate dehydrogenase activity |
|  |  |  | 4.90E-10 | 14 | 69 | 12.3 | Protein domain specific binding |
|  |  |  | 5.70E-08 | 13 | 82 | 9.6 | NAD binding |
|  |  | GO Biological Process | 4.40E-38 | 56 | 317 | 10.7 | Response to cadmium ion |
|  |  |  | 9.90E-34 | 59 | 433 | 8.2 | Response to metal ion |
|  |  |  | 1.00E-47 | 104 | 1055 | 6 | Carboxylic acid metabolic proc. |

### Identification of proteins with significant change in abundance (DAPs)

Quantitative differences occurring among proteome profiles due to the different treatments were detected by comparing the individual protein intensities in each treatment group (Stress, Inhibitor+Stress and Inhibitor) with the control samples. Protein abundance differences were obtained by performing a t-test (p<0.05) on the proteins shown to be significant different in any of the treatments, i.e., 581 proteins after 10 min of treatment and 909 proteins after 30 min of treatment (indicated in Fig.2A) and visualized in volcano plots (Fig. 4A and B). This analysis resulted in 49 DAPs (18 more abundant and 31 less abundant) between Stress treatment and control after 10 min and 65 DAPs (22 more abundant, 43 less abundant) after 30 min. For the Inhibitor only treatment a total of 48 DAPs (7 higher abundant, 41 less abundant) were found after 10 min and 84 DAPs (26 more abundant, 58 less abundant) in the 30 min set. The highest number of differences occurred for the Inhibitor+Stress treatment versus control, with 459 DAPs (287 more abundant, 172 less abundant) after 10 min and 667 DAPs (468 more abundant, 199 less abundant) after 30 min. The DAPs identified for each treatment were further subjected to comparable analysis to categorise them based on whether they require Ca^2+^ for their regulation.

**Fig. 4:**
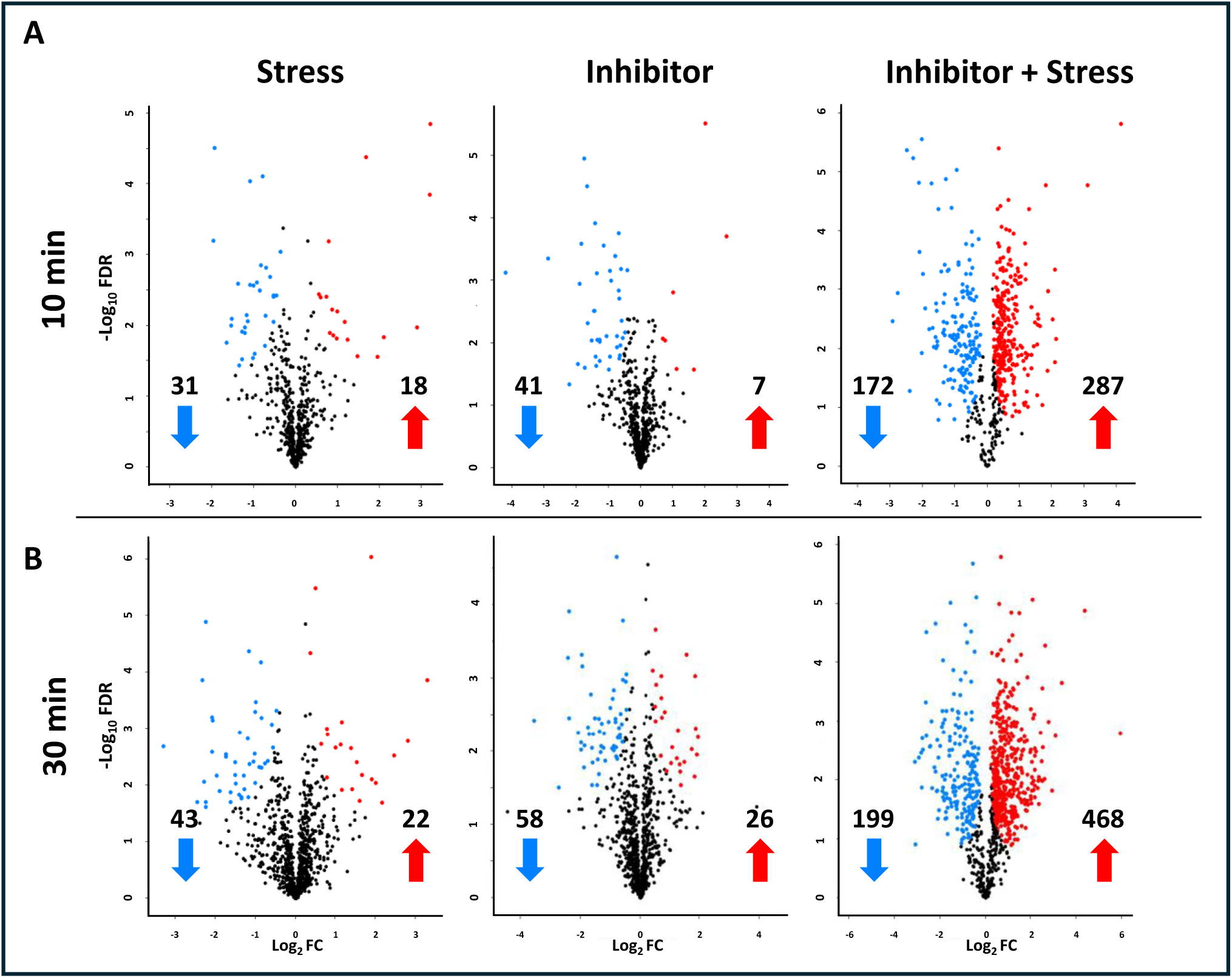
Volcano plots indicating DAPs for three treatments (Stress, Inhibitor and Inhibitor+Stress) in comparison to control (double mock) samples (FDR 0.05) after 10 ’min (**A)** and 30 min **(B)** of treatment. Proteins were graphed by fold change (x-axis) and the confidence statistic (-log P) on the y-axis. Blue dots represent proteins that show a significant lower abundancy while red dots represent proteins that show a significant higher abundancy for the indicated treatment compared to control samples. Black dots represent the proteins that do not show a significant change in abundance (S0=0.1, FDR= 0.05).

### Comparison of DAPs and identification of Ca^2+^ dependent and independent proteins

After identification of the DAPs for each treatment vs control at both timepoints (Fig. 4), the DAPs of each treatment were compared to all other treatments of the same timepoint to find overlapping and unique proteins (Fig. 5A). In general, the overlapping numbers were relatively low, indicating that each treatment has an individual effect on the proteome. For both time points the largest overlap was observed between Stress and Inhibitor+Stress as well as Inhibitor and Inhibitor+Stress with a much smaller overlap between Stress and Inhibitor. As the total number of proteins differs between the two time points, we decided to also compare the relative numbers (% from total proteins) of DAPs (suppl. Fig. 1). This comparison shows a very similar pattern between the two time points.

**Fig. 5.**
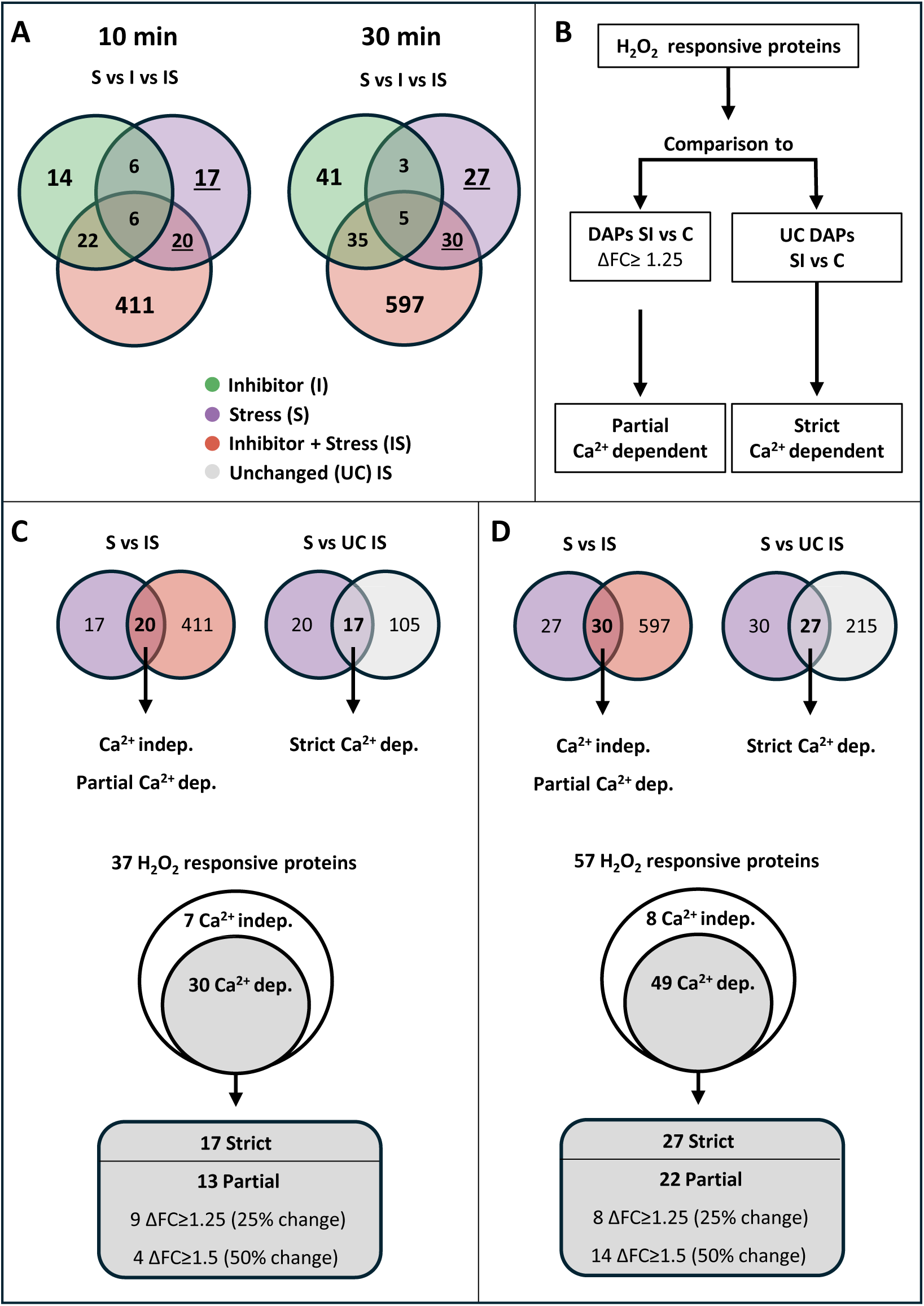
Identification and categorisation of H_2_O_2_ responsive proteins. **A:** Venn diagrams showing all identified DAPs and their overlap between the different treatments. Underlined numbers are the H_2_O_2_ responsive proteins. **B:** Schematic representation of the analysis steps to identify the different levels of Ca^2+^ dependency for H_2_O_2_ responsive proteins. **C:** Categorisation of the H_2_O_2_ responsive proteins after 10 min of stress treatment. **D:** Categorisation of the H_2_O_2_ responsive proteins after 30 min of stress treatment

For the further analysis, we wanted to focus on H_2_O_2_ responsive proteins, i.e. those proteins that show a difference in abundance between H_2_O_2_ treatment and control (DAP-Stress, 10 and 30 min). From these sets we omitted the DAPs that showed a different abundance upon treatment with LaCl_3_ alone, to avoid effects of the inhibitor that are not related to the reduction of the H_2_O_2_ induced Ca^2+^ transient. This resulted in a set of 37 H_2_O_2_ responsive proteins after 10 min of treatment and 57 proteins after 30 min of treatment (Fig. 5A, underlined numbers). These stress responsive proteins were further categorised as being Ca^2+^-independent or Ca^2+^-dependent, depending on their abundance in the Inhibitor+Stress treatment. For this categorisation we used the method described by Bhattacharyya et al. (2025). The steps used in this approach are schematically displayed in Fig. 5B. H_2_O_2_ responsive proteins that showed an unchanged abundance (UCs) compared to control under Inhibitor+Stress treatment can be considered strictly Ca^2+^ dependent in their H_2_O_2_ response (Fig. 5C and D). Proteins that showed a differential abundance upon both Stress vs. control and Stress+Inhibitor vs. control, but their abundance level differed significantly (ΔFC ≥1.25, corresponding to a change in protein abundance of at least 25%) between the two treatments, were categorised as being partially Ca^2+^ dependent (Fig. 5C and D). This group was further split in two categories, with the lower threshold set to a change in abundance of at least 25% up to 50%, and the higher threshold including all proteins showing a change in abundance of at least 50% (ΔFC ≥1.5). Proteins that showed no differential abundance between Stress and Stress+Inhibitor treatment (ΔFC ≤1.25) were considered Ca^2+^- independent in their H_2_O_2_ response.

This analysis identified a total of 7 Ca^2+^-independent and 30 Ca^2+^-dependent H_2_O_2_ responsive proteins after 10 min of treatment (Fig. 5C). Among the Ca^2+^-dependent proteins, 17 were classified as strictly Ca^2+^-dependent and 13 partial Ca^2+^-dependent. Of these 13 partial Ca^2+^-dependent proteins, 4 met the threshold of at least a 50% change in abundance (≥2-fold increase or decrease). For the 30 min treatment, 8 Ca^2+^-independent and 49 Ca^2+^-dependent H_2_O_2_ responsive proteins were identified (Fig. 5D). Within the Ca^2+^-dependent group, 27 proteins were strictly Ca^2+^-dependent, a similar percentage as observed at 10 min, while 22 proteins displayed partial Ca^2+^-dependence. Among the partially Ca^2+^- dependent proteins, 14 met the threshold of at least a 50% change in abundance (≥2-fold increase or decrease). A detailed list of the H_2_O_2_ responsive proteins, along with their classification as Ca^2+^- independent, partially Ca^2+^-dependent or strictly Ca^2+^-dependent, is provided in Table 3 for the 10 min and Table 4 for the 30 min data set.

**Table 3:** List of H_2_O_2_ responsive proteins indicated as strict Ca^2+^ dependent (dark grey), partial Ca^2+^ dependent (light grey) and Ca^2+^ independent (white) after 10 min of treatment. Partial Ca^2+^ dependent proteins were further divided by a threshold of at least a 50% change in abundance (≥2-fold increase or decrease).

| Protein ID |  |  | Abundance | Full name | ΔFC | Symbol |
| --- | --- | --- | --- | --- | --- | --- |
|  |  |  | (compared to control) |  | (SvsC-SlvsC) |  |
| Strict Ca<sup>2+</sup> dependent |  | AT1G16460 | Less | Sulfurtransferase 2 |  | STR2 |
|  |  | AT1G48350 | More | Large ribosomal subunit protein uL18c |  | RPL18 |
|  |  | AT1G62180 | Less | 5' adenylylsulfate reductase 2, chloroplastic |  | APR2 |
|  |  | AT1G72610 | More | Germin like protein |  | GLP1 |
|  |  | AT2G30520 | More | Root phototropism protein 2 |  | RPT2 |
|  |  | AT2G44060 | Less | Late embryogenesis abundant protein |  | LEA26 |
|  |  | AT3G07660 | Less | Flocculation protein |  | DUF1296 |
|  |  | AT3G14990 | Less | Protein DJ-1 homolog A |  | DJ1A |
|  |  | AT3G17880 | Less | TPR repeat- containing thioredoxin |  | TDX |
|  |  | AT3G17900 | Less | Heat-inducible transcription repressor |  | MEB5_12 |
|  |  | AT3G45850 | Less | Kinesin motor domain- containg protein |  | KIN5D |
|  |  | AT3G53260 | Less | Phenylalanine ammonia-lyase 2 |  | PAL2 |
|  |  | AT4G28730 | Less | Glutaredoxin-C5 chloroplastic |  | GRXC5 |
|  |  | AT5G06460 | Less | Ubiquitin-activating enzyme |  | UBA2 |
|  |  | AT5G21274 | Less | Calmodulin |  | CAM |
|  |  | AT5G43830 | Less | Domain containing protein |  | DUF3700 |
|  |  | AT5G59890 | Less | Actin-depolymerizing factor 4 |  | ADF4 |
| Partial Ca<sup>2+</sup> dependent | ΔFC ≥ 1.5 | AT1G13750 | More | Probable inactive purple acid phosphatase 1 | 5.37 | PAP1 |
|  |  | AT1G20160 | More | CO(2)-response secreted protease | 2.51 | CRSP |
|  |  | AT3G49080 | Less | Small ribosomal subunit protein uS9m | 1.71 | RPS9M |
|  |  | AT4G38510 | More | V-type proton ATPase subunit B2 | 2.34 | VHA-B2 |
|  | ΔFC 1.25 - 1.5 | AT1G10200 | Less | LIM domain-containing protein | 1.26 | WLIM1 |
|  |  | AT1G14030 | More | Fructose-bisphosphate aldolase | 1.39 | LSMT-L |
|  |  | AT1G26460 | More | Tetratricopeptide repeat-containing protein | 1.32 | TPR |
|  |  | AT1G70890 | Less | MLP-like protein 43 | 1.45 | MLP43 |
|  |  | AT2G20890 | More | Thylakoid formation 1 | 1.29 | THF1 |
|  |  | AT3G53130 | More | Carotene epsilon-monooxygenase, chloroplastic | 1.29 | CYP97C1 |
|  |  | AT4G11260 | Less | SGT1 homolog B | 1.40 | SGT1B |
|  |  | AT4G14440 | More | Enoyl-CoA delta isomerase 3 | 1.28 | ECI3 |
|  |  | AT4G29510 | Less | Protein arginine N-methyltransferase 1.1 | 1.34 | PRMT11 |
| Ca<sup>2+</sup> independent | ΔFC <1.25 | AT1G11790 | Less | Arogenate dehydratase | 1.16 | ADT1 |
|  |  | AT2G20900 | More | Diacylglycerol kinase 5 | 1.11 | DGK5 |
|  |  | AT2G42690 | More | Phospholipase A1-IIdelta | 1.07 | AGAP1 |
|  |  | AT3G45140 | More | Lipoxygenase 2, chloroplastic | 1.10 | LOX2 |
|  |  | AT4G13010 | More | Chloroplast envelope quinone oxidoreductase homolog | 1.13 | CEQORH |
|  |  | AT4G28660 | More | Photosystem II reaction center Psb28 protein | 1.00 | PSB28 |
|  |  | AT5G63870 | Less | Serine/threonine-protein phosphatase 7 | 1.09 | PP7 |

**Table 4:**
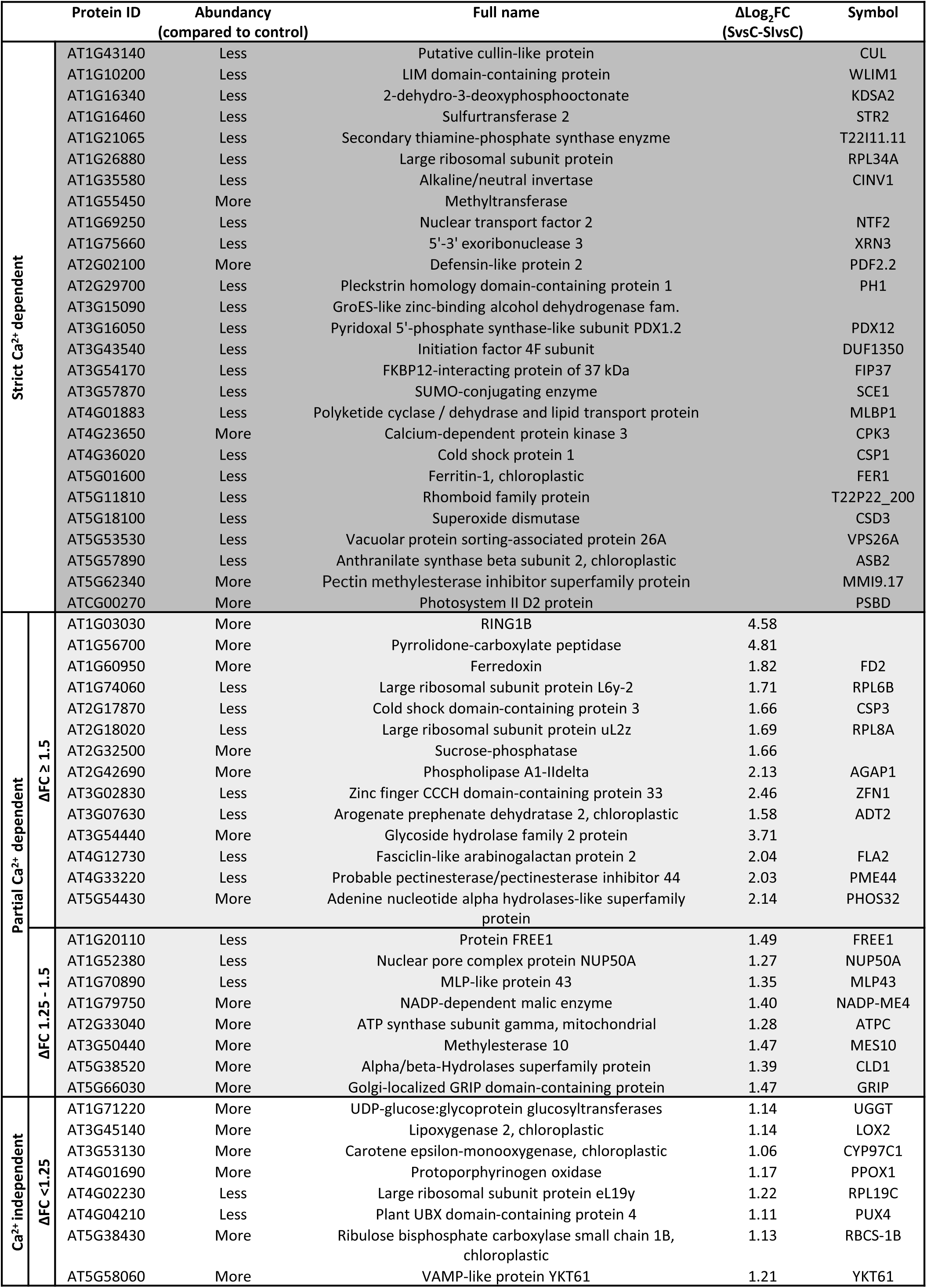
List of H_2_O_2_ responsive proteins indicated as strict Ca^2+^ dependent (dark grey), partial Ca^2+^ dependent (light grey) and Ca^2+^ independent (white) regulated after 30 min of treatment. Partial Ca^2+^ dependent proteins were further divided by a threshold of at least a 50% change in abundance (≥2-fold increase or decrease).

### Effect of the duration of the stress treatment

As stated above, the largest difference in biological function was observed between the two time points of stress treatment. To further elucidate the effects of stress duration on DAPs, a comparative analysis was conducted using two approaches. First, the absolute numbers of H_2_O_2_ responsive proteins identified after 10 and 30 min of stress treatment were compared. This comparison, visualized in a Venn diagram (Fig. 7A, upper panel), revealed a duration related increase of H_2_O_2_ responsive proteins from 37 to 57, with an overlap of only six proteins between the two time points. The latter is in line with the different biological processes observed for the proteins in the two data sets (Fig. 6A and B) and indicates that the duration of the treatment results in a significant different effect on the proteome. The same trend holds true, when proteins were separated based on their Ca^2+^ dependency (Fig. 7A, lower panel).

**Fig. 6:**
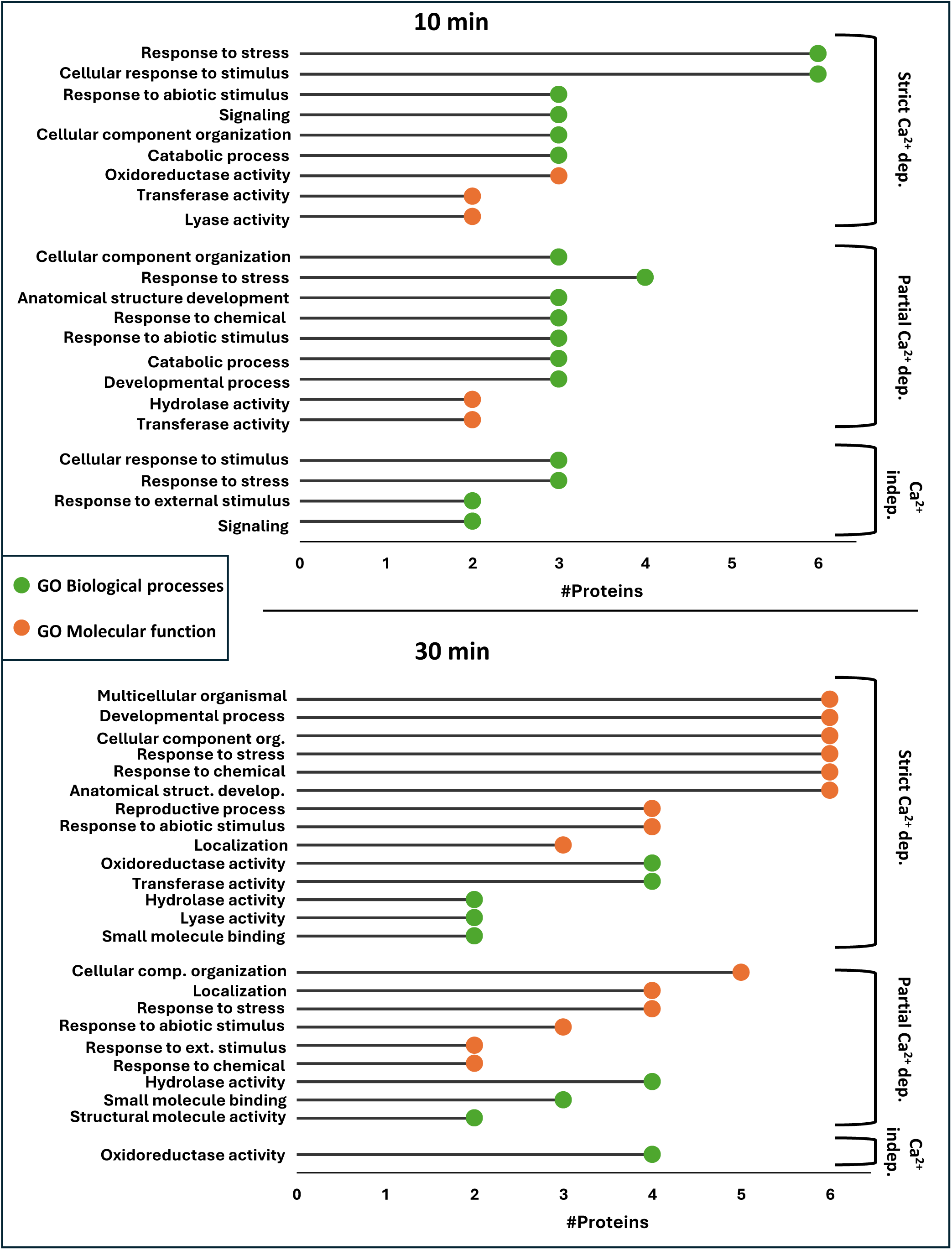
GO-term analysis (Biological Processes and Molecular Function) on the H_2_O_2_ responsive proteins indicated as strict Ca_2+_ dependent, partial Ca_2+_ dependent, and Ca_2+_ independent regulated after 10 and 30 min of treatment. Numbers represent absolute numbers of proteins that are annotated to functional categories based on high level GO terms.

**Fig. 7:**
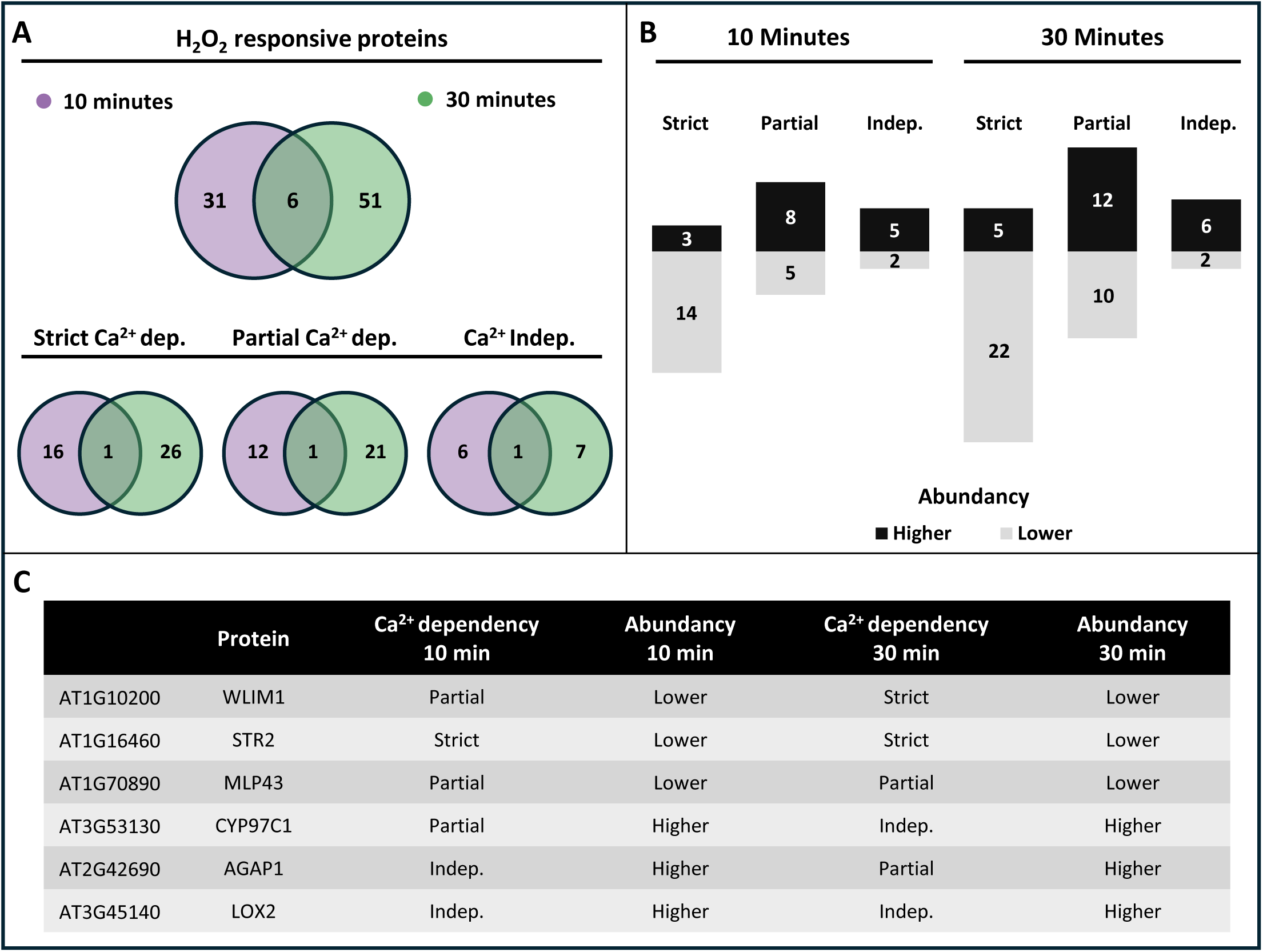
Comparison of the H_2_O_2_ responsive proteins between 10 and 30 min of stress treatment. **A:** Venn diagrams for comparison of all H_2_O_2_ responsive proteins identified after 10 min (purple) and 30 min (green) of stress treatment (top part). Comparison of the H_2_O_2_ responsive proteins after 10 and 30 min of stress treatment, separated for their dependency on Ca^2+^ (bottom). Values represent absolute number of DAPs in each category. **B**: Bar chart showing absolute numbers of DAPs and their change in abundancy after H_2_O_2_ treatment separated for the group of Ca^2+^ dependency they resolve into. **C**: Overview of the Ca^2+^ dependency and the change in abundancy after stress treatment of the 6 H_2_O_2_ responsive proteins found to be overlapping between 10 and 30 min of stress treatment.

We further investigate the effect of stress duration by additionally considering the increase and decrease of proteins abundance (Fig. 7B). Although the total number of DAPs was relatively low, a clear pattern emerged. Proteins that were strictly Ca^2+^-dependent predominantly exhibited a significant decrease in abundance, whereas those that were partially Ca^2+^-dependent or Ca^2+^-independent tended to show increased abundance.

We than had a closer look at the only six H_2_O_2_ responsive proteins that were found at both timepoints (Fig. 6A, upper panel). Their Ca^2+^ dependency and changes in abundance were examined across both conditions (Fig. 6C). The analysis revealed that all six proteins exhibited the same change in abundance (either increased or decreased) after both 10 and 30 min of stress treatment. Three of these proteins - WLIM1, CYP97C1, and AGAP1 - displayed a difference in their Ca^2+^ dependency between the two timepoints.

### Validation of the dataset

We tried to assess the accuracy of the Mass spec analysis by comparing the data output (raw average LFQ intensity values) with Western blot analysis for two proteins from our dataset, for which antibodies could be obtained (Fig. 8). PHENYLALANINE LYASE 2 (PAL2), known to be responsive to oxidative stress (Stanley Kim et al., 2005), showed a lower abundance after 10 min of H_2_O_2_ treatment (Fig. 8A, upper panel) and was found in the group of strict Ca^2+^-dependent H_2_O_2_ responsive proteins (Table 3). This pattern could be confirmed by the Western blot analysis, where the protein band detected in the extracts of stress treated plant material was clearly fainter as for the other treatments (Fig.8A, lower panel). The aquaporin GAMMA TONOPLAST INTRINSIC PROTEIN 2 (TIP2), suspected to be involved in hydrogen peroxide transmembrane transport (Bienert et al., 2007), showed a higher abundance in the 10 min stress treated samples compared to control but is not in Table 4 due to its also higher LFQ-values after inhibitor only treatment (Fig. 8B upper panel). Also, this result could be confirmed by the increased intensity of the reaction in the stress treated and the inhibitor treated samples in the Western blot (Fig.8B, lower panel). Thus, in both cases an agreement between the proteome data (LFQ values) and the Western blot analysis was found.

**Fig. 8:**
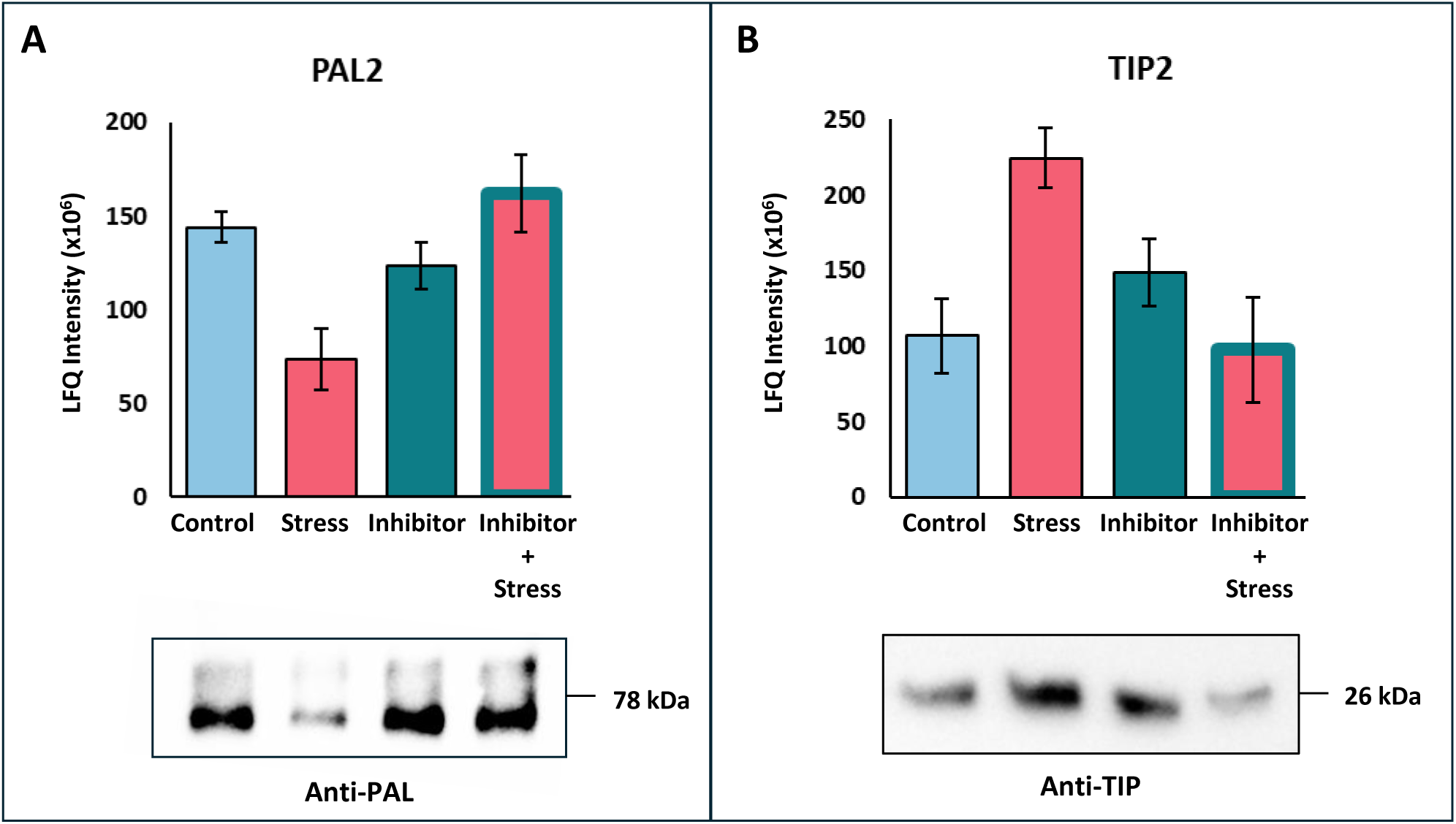
Western blot validation of proteomic data. Averages (n=5) of the raw LFQ intensity values (bar graph) and immunodetection using specific antibodies for (blot) for **A:** PAL2 and **B:** TIP2. Full sized blots are shown in Supplementary Fig. S2

## Discussion

In this study, we employed a pre-treatment with the Ca^2+^ channel inhibitor La^3+^ to differentiate between Ca^2+^-dependent and Ca^2+^-independent changes in protein abundance after treatment with H_2_O_2_ in *Arabidopsis thaliana* at a proteome-wide scale. Our investigation was focused on short-term responses, therefore we analysed the proteome after 10 and 30 min of treatment with H_2_0_2_. We detected 3724 proteins after 10 min and 3757 after 30 min of treatment. From these, 581 and 909 proteins significantly changed abundance, respectively. The much smaller number of proteins used for further analysis is the result of proteomics experiments often generating a large number of low-confidence identifications or proteins with inconsistent quantifications across replicates.

A key initial step in our analysis was the identification of H_2_O_2_-responsive proteins, which resulted in distinct subsets of proteins detected after 10 and 30 min of stress treatment, with only six proteins identified in both sets. This temporal variation in protein abundance highlights the dynamic nature of the oxidative stress response and suggests that the duration of stress exposure significantly influences proteome-wide adaptations. Given that H_2_O_2_ is known to induce Ca^2+^signals at the cellular level (Rentel & Knight, 2004), which are then decoded by Ca^2+^-binding proteins to activate downstream molecular processes (Mohanta et al., 2019; Tang et al., 2020), the observed temporal differences are in line with the described spatiotemporal plasticity of Ca^2+^signaling (Boulware & Marchant, 2008). Consequently, it is not unexpected that a threefold increase in stress duration results in distinct proteomic responses, reflecting the dynamic and evolving nature of oxidative stress adaptation at the molecular level.

Following the identification of H_2_O_2_-responsive proteins, we further categorized them based on their dependence on Ca^2+^ for differential abundance regulation. Strict Ca^2+^ dependency was defined by proteins that exhibited significant changes in abundance upon H_2_O_2_ treatment but lack this response when pre-incubated with LaCl_3_, indicating a complete reliance on Ca^2+^ signaling for their regulation. This was the largest group for both the 10 and 30min time point. Partially Ca^2+^-dependent proteins showed a difference in abundance both between control and H_2_O_2_ treatment as well as control and H_2_O_2_+LaCl_3_ treatment, however, the abundance was significantly different between the two treatments. Ca^2+^ independency was observed for less than 20% and 15% of the H_2_O_2_-responsive after 10 and 30 min, respectively, showing the strong impact of Ca^2+^ signaling on the oxidative stress response that was also observed in a recent transcriptomic analysis on barley (Bhattacharyya et al., 2025). Another notable finding related to this was the high number of DAPs identified when the H_2_O_2_- induced Ca^2+^ transient was blocked by LaCl_3_. With over 400 DAPs at 10 and over 600 DAPs at 30 min, the numbers were about a factor 10 higher than for the stress treatment or inhibitor treatment alone. The high number of DAPs in response to the combined treatment suggest that Ca^2+^ signaling can strongly attenuate the H_2_O_2_ response.

Another interesting result is the observation that much more proteins with a strict Ca^2+^ dependency show a reduced abundance, while a higher abundance is more often observed for Ca^2+^ independent H_2_O_2_-responsive proteins. Considering the timeframe of the experiment it seems likely that protein loss is mostly driven by degradation (and not reduced transcription) while increase in protein content is the result of increased transcription and/or translation. These findings indicate a potential regulatory mechanism in which protein destabilization is driven by Ca^2+^ signaling, while proteins without strict Ca^2+^ dependency may undergo enhanced synthesis under oxidative stress conditions.

Comparison of H_2_O_2_-responsive proteins between 10 and 30 min of treatment revealed only six overlapping proteins, all of which exhibited the same change in abundance at both time points. Notably, three proteins exhibited a shift in Ca^2+^ dependency between the two time points. However, they went from partial to strictly Ca^2+^ dependent or from partial to Ca^2+^ independent, but no protein changed from strictly Ca^2+^-dependent to Ca^2+^-independent. Since the difference in logFC change between the two time points is quite consistent for these three proteins over the five biological replicates analysed, these findings clearly suggest a dynamic nature of Ca^2+^ dependent and independent protein regulation in response to oxidative stress.

Overall, it remains challenging to draw a definitive conclusion regarding the precise role of Ca^2+^ signaling in shaping proteomic changes from our data. However, we found several candidates with known functions in stress responses among the H_2_O_2_-responsive proteins. In all of these cases single proteins and not proteins groups were identified. The ribosomal protein RPL18, which exhibited higher abundance after 10 min of H_2_O_2_ treatment, has been previously described as a positive regulator of powdery mildew resistance in wheat (Tao et al., 2024). The germin-like protein GLP1, which also showed increased abundance after 10 min, has been characterized as an oxidative stress defence enzyme in plants (Shahwar et al., 2023). CPK3, a calcium-dependent protein kinase, displayed higher abundance with strict Ca^2+^ dependency after 30 min, aligning with its known role in Ca^2+^-dependent signaling pathways involved in responses to abiotic and biotic stresses (Mehlmer et al., 2010) and plant immunity (Lu et al., 2020). For the identified H_2_O_2_-responsive proteins that lack a well-defined role in plant stress responses, further investigation is required to elucidate their functions in plant defence mechanisms, signaling pathways, ROS homeostasis, and overall plant survival.

### Conclusion

Hydrogen peroxide (H_2_O_2_) is a crucial reactive oxygen species (ROS), generated as a toxic by-product of biological metabolic processes while also functioning as a signaling molecule that regulates plant growth and development. Additionally, it interacts with signaling pathways involving second messengers such as Ca^2+^. Our findings expand the current knowledge of oxidative stress responses by identifying proteins which are degraded or synthesized in response to H_2_O_2_ in a Ca^2+^ dependent manner. In these subsets, proteins were found that are known to play a role in Ca^2+^ signaling and stress response, but also proteins that were unassociated with stress response pathways before. These novel proteins present potential targets for further investigation into the molecular basis of H_2_O_2_-Ca^2+^ interactions. Given that both biotic and abiotic stress factors can induce H_2_O_2_ accumulation and Ca^2+^ fluctuations, understanding this crosstalk is essential for deciphering plant stress acclimation mechanisms. The insights gained from Arabidopsis may thus have broader applications, potentially for strategies to enhance stress resilience in economically significant crop species.

## Material and Methods

### Plant material and growth conditions

Leaf protein extracts for the proteomics analysis were obtained from *A. thaliana* (ecotype Columbia; Col-0). Seeds were sown on soil, stratified for 2 days at 4°C in the dark, and separated after germination into single pots filled with standard plant potting soil pre-treated with Confidor WG 70 (Bayer Agrar, Germany). Plants were cultivated in a climatized growth chamber with a room temperature of 20 ± 2°C, a light intensity of ∼150 µmol photons m^-2^ s^-1^ (Philips TLD 18 W of alternating 830/840 light colour) and long day conditions (16 h light/8 h dark). Pre-experiments to determine the best conditions for the inhibitor treatment and stress stimulus were performed with Arabidopsis plants expressing cytosolic apoaequorin (At-AEQ_cyt_) (M. R. Knight et al., 1991).

### Aequorin reconstitution, luminescence measurements and Ca^2+^ concentration calculations

Stimulus-induced Ca^2+^ transients were analysed using leaf material of three-week-old At-AEQ_cyt_ plants. The day before measurements were taken, leaf discs (Ø 6mm) were collected and incubated overnight in the dark at 20°C in 5 µM coelenterazine (Biosynth AG, Switzerland) for reconstitution of the cytosol targeted apoaequorin to aequorin. After reconstitution, leaf discs were carefully washed (ddH_2_O) and transferred either into 1 mM LaCl_3_ solution (inhibitor pre-treatment) or ddH_2_O (mock) for 1 hour. Subsequently, single leaf discs were washed again and transferred individually into a 96-well plate (Lumitrac 600, Greiner Bio-One, Austria), floating in 100 µl ddH_2_O. Photon count measurements were performed using a plate luminometer (Tristar 2 Multimode Reader, Berthold GmbH). First the basal level of photon counts was measured for 30 seconds with an interval of 1 second, followed by the application 20 mM H_2_O_2_ using a 40 mM stock solution and a volume equal to the starting volume of the ddH_2_O (100 µl), with continuous measuring of the response for 240 seconds. The remaining aequorin was discharged by adding discharge solution (final concentration of 1 M CaCl_2_ in 10% (v/v) EtOH) and photon counts were recorded for another 300 sec. Concentrations of free calcium ions in the cytosol ([Ca^2+^]_cyt_) were calculated based on the photon counts as described before (H. Knight & Knight, 1995). The measurements were performed with three independent experimental replicates consisting of three technical replicates.

### Sample collection and treatment for proteomics

For proteomics analysis, 12 complete rosettes of three-weeks-old Col-0 plants were incubated in 1 mM LaCl_3_ (inhibitor treatment) or ddH_2_O for 1 hour. ddH_2_O and LaCl_3_ pre-treated plants were carefully washed and then transferred into either 20 mM H_2_O_2_ (stress treatment) or ddH_2_O for the control treatment. For each proteomics sample, complete rosettes of 12 plants were harvested after 10 and 30 min of the stress treatment, pooled, immediately frozen in liquid nitrogen, and stored at −80 °C until protein extraction. Within one experiment, plants from all four treatments were harvested for both timepoints (10 and 30 min), and a total of 5 experiments with independently grown plants were performed. For a schematic overview of the protocol see Figure 1.

### Protein isolation, precipitation, lysis and digestion

Frozen plant material was first ground in liquid nitrogen using a pre-cooled mortar and pestle. 500 mg of the ground plant material was mixed with 2 ml ice-cold Lacus protein isolation buffer (20 mM Tris [pH 7.7], 80 mM NaCl, 0.75 mM EDTA, 1 mM CaCl_2_, 5 mM MgCl_2_, 1 mM DTT, 1 mM NaF) containing 4 tablets of protease inhibitor (Roche cOmplete EDTA-free, protease inhibitor cocktail tablets) and 10 tablets of phosphatase inhibitor (Roche PhosSTOP) per 200 ml. Samples were incubated on ice for 10 min followed by centrifugation at 15000 g for 10 min at 4 °C. Supernatants were transferred into a fresh tube. An equal volume of 20 % (w/v) trichloroacetic acid (TCA) was added to the supernatant, and the samples were placed on ice for 30 min. Afterwards, the samples were centrifuged at 15000 g for 10 min at 4 °C and the supernatant removed. The precipitated protein pellets were washed with cold 80% acetone and the samples were vacuum-dried. 50 µl of urea lysis buffer (8 M urea, 150 mM NaCl, 40 mM Tris-HCl pH 8) was added and the protein concentration was determined via the Pierce^TM^ BCA Protein Assay (Thermo Fisher Scientific). Subsequently, 3 mg total protein per sample was reduced in 5 mM DTT and alkylated in 15 mM iodoacetamide for 30 min in the dark at room temperature. The alkylated samples were quenched by adding DTT to a final concentration of 5 mM and mixed with 30 mg carboxylate beads (Sera-Mag^TM^, 1:1 ratio of hydrophilic and hydrophobic beads, Cytiva, USA). Proteins attached to the beads were washed four times with 80 % (v/v) ethanol and digested in ammonium bicarbonate buffer (30 mM, pH 8.2) containing 30 µg Trypsin (Promega, WI, USA). Tryptic digestion was performed overnight at 37 °C under constant shaking. The digestion was stopped by the addition of formic acid (final concentration 4%). 100 µg of the digested peptides per sample were transferred into a new reaction tube, vacuum-dried and stored at −20 °C until HPLC-MS analysis.

### LC-MS analysis

Digested peptides were subjected to LC-MS analysis at the Mass Spectrometry unit of the faculty of life sciences at the University of Vienna as described previously (Bleker et al., 2024). In brief: approximately 1 µg of peptide sample was reconstituted in 0.1% (v/v) formic acid and separated using an online reversed-phase high-pressure liquid chromatography (HPLC) system. Separation was performed on a heated C18 analytical column over a 140-minute gradient (5-50%). The eluate was introduced into a Q-Exactive Plus mass spectrometer using an Easy-Spray ion source. Mass spectra were acquired in positive ion mode with a data-dependent acquisition strategy, selecting the top 15 most intense ions for MS analysis. A full MS scan was performed at 70,000 resolution (m/z 200), followed by MS/MS fragmentation at 17,500 resolution using higher-energy collisional dissociation (HCD) at 27% normalized collision energy. Dynamic exclusion was set to 40 seconds, and specific precursor ions (unassigned, +1, +7, +8, and >+8 charge states) were excluded. The analysis was conducted with five independent experimental replicates per sample to ensure reproducibility.

### Peptide identification and quantification

Identities and peptide features were defined by the peptide search engine Andromeda provided by the MaxQuant software (Prianichnikov et al., 2020) and using standard settings (Tyanova et al., 2016). In detail: trypsin-based digestion of the peptides with up to two missing cleavage sites was selected. Methionine oxidation as well as N-terminal acetylation was set as a variable modification for peptide identification. In total, up to three potential modification sites per peptide were accepted. The identified peptide sequences were searched and aligned against the Araport11 reference protein database (Cheng et al., 2017). The false discovery rate cut-off for protein identification and side identification was set to 0.01. The minimum peptide length was set to seven and the maximum length to 40 amino acids. For each identified protein group, label-free quantitation (LFQ) intensities were calculated using the Maxquant software. A protein group contains all proteins and protein isoforms that cannot be unambiguously identified by unique peptides but have shared peptides. We further on refer to protein groups only as ‘proteins’.

### Data analysis

For quantitative proteome analyses, the derived LFQ intensities were loaded into the Perseus software (Tyanova et al., 2016) and used for data and statistical analysis, as well as graphics and visualisation of the results. The steps taken to determine which proteins show a differential abundance among the different treatments was based on a method described before (Nikonorova et al., 2018). In short: after loading the LFQ intensities, a quality control was performed in which protein groups with the indication ‘only identified by site’ (proteins that are only identified by peptides carrying modified amino acids), reverse sequences (decoy proteins), and potential contaminants (for example, albumin) were filtered out. Biological replicates were grouped, and values were *log2* transformed. Protein groups were filtered based on valid values, where the criterium was set to have at least 3 valid values in one group (each treatment group consists of 5 replicates) to remove the low abundant proteins. Remaining missing values (protein group not identified in a run) were imputed with values based on the normal distribution with a width of 0.3 (relative to the standard deviation of the measured values) and a downshift of 1.8. The imputed numbers represent very small values, meaning the identified peptide has a very low abundance. Principal component analysis (PCA) was performed using the Perseus software. Differential abundant proteins were determined by multiple sample ANOVA test (p-value < 0.05). P-values were corrected for multiple testing using Benjamin-Hochberg rule (adjusted P-value). All ANOVA significant proteins were z-score normalized and used for supervised hierarchical clustering to produce a heatmap, using Euclidean distance and average linkage. Pairwise Student’s t-tests (p-value < 0.05) were performed on the non-z-scored values to determine differences in protein abundance between two treatments. Volcano plots were generated using the Perseus software, by plotting log_2_ fold-change values on the x-axis against the -log p values on the y-axis, cut-off was set by nonlinear volcano lines based on S0 = 0.1 adjusted p-value. Protein groups showing different abundance among the treatments were analysed for overlapping groups between treatments and time points. Gene ontology (GO) enrichment analysis (KEGG, biological processes, molecular function) was performed on the major protein clusters identified by hierarchical clustering, and on the identified H_2_O_2_ responsive proteins using ShinyGO, which uses the annotations of Ensembl and STRING-db (Ge et al., 2020). An error probability according to Fisher’s’ t-test of <0.05 and a false-discovery rate (FDR) of <0.01 was selected for enriched GO-Terms.

### Immunodetection

For immunodetection, proteins were isolated from the plant material that was used for the MS-analysis using the same Lacus protein isolation buffer and isolation protocol (described above). The proteins were separated on a 12% SDS-polyacrylamide gel and transferred to nitrocellulose membrane (0.45µm pore size; Bio-Rad Laboratories). After transfer of the proteins, the membrane was stained using 0.1% (w/v) Ponceau S in 5% (v/v) glacial acetic acid. Immunodetection was performed using antibodies against Anti-PAL 1-4 (dilution 1:2000) and Anti-TIP 1;1-2 (dilution 1:1000) (Agrisera, AS214614 and AS22 4844). Blots were incubated with the matching secondary antibody (anti-rabbit IgG horse radish peroxidase conjugated, AS09602, dilution 1:25000) and developed with the

### Data availability statements

The authors upon reasonable request can provide the data presented in this study

## Author contributions

A.v.D. contributed to conceptualization, investigation, formal analysis, validation, visualization, and writing - original draft as well as review & editing; A.B. and B.W. contributed to conceptualization and sample preparation for LC-MS measurements; L.A. contributed to LC-MS analysis and writing the corresponding method; W.W contributed to supervision; M.T. contributed to conceptualization, supervision and writing – review editing; U.C.V. contributed to conceptualization, formal analysis, validation, funding acquisition, project administration, supervision, and writing – review & editing.

