## Supplemental Material for "With or without a Ca^2+^ signal? A proteomics approach towards Ca^2+^ dependent and independent proteome changes in response to oxidative stress in *A. thaliana*"

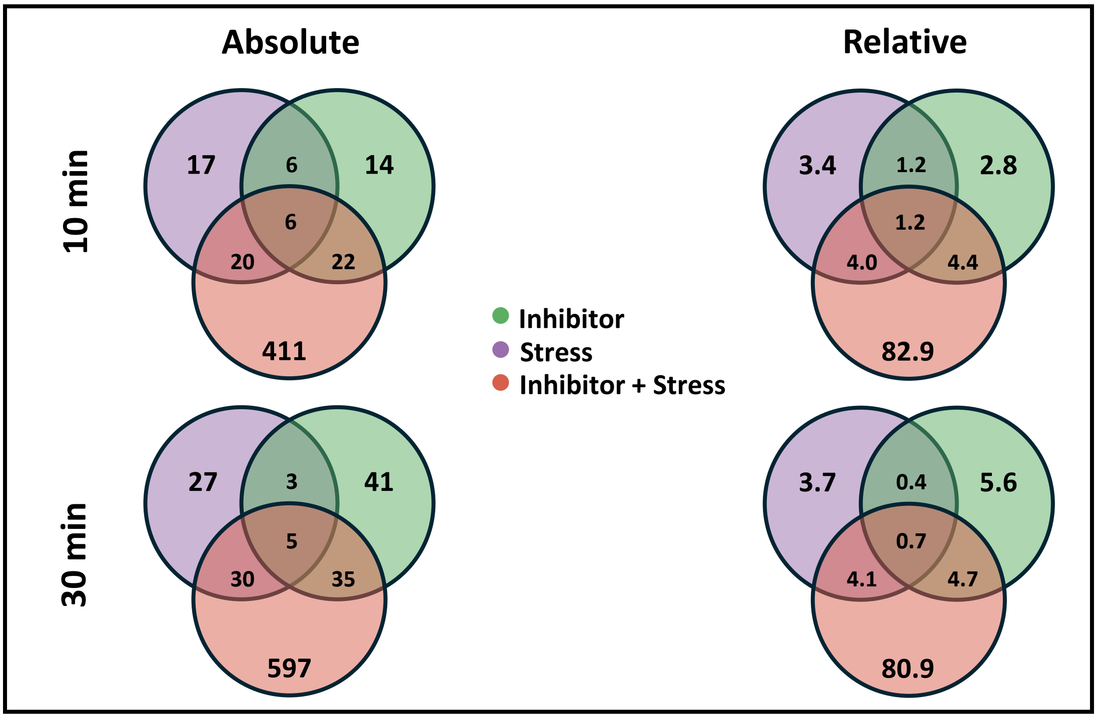


**Supplementary Fig. S1:** Absolute and relative (% of total identified DAPs) number of DAPs for 10 min (top) and 30 min (bottom) of stress treatment.

| | Pathway database (GO) | GO Functional category | # Proteins | Proteins |
|---|---|---|---|---|
| Strict $Ca^{2+}$ dependent | Biological process | Cellular response to stimulus | 6 | RPT2 DJ1A GRXC5 UBA2 CAM6 ADF4 |
| | Biological process | Response to stress | 6 | DJ1A TDX PAL2 GRXC5 UBA2 ADF4 |
| | Biological process | Response to abiotic stimulus | 4 | RPT2 TDX PAL2 |
| | Biological process | Catabolic process | 3 | DJ1A PAL2 UBA2 |
| | Biological process | Cellular component organization | 3 | TDX GRXC5 ADF4 |
| | Biological process | Signaling | 3 | RPT2 CAM6 ADF4 |
| | Molecular function | Oxidoreductase activity | 3 | APR2 TDX GRXC5 |
| | Molecular function | Transferase activity | 2 | STR2 UBA2 |
| | Molecular function | Lyase activity | 2 | DJ1A PAL2 |
| Partial $Ca^{2+}$ dependent | Biological process | Response to stress | 4 | CRSP MLP43 THF1 SGT1B |
| | Biological process | Developmental process | 3 | CRSP RPS9M SGT1B |
| | Biological process | Catabolic process | 3 | THF1 SGT1B ECI3 |
| | Biological process | Response to abiotic stimulus | 3 | THF1 SGT1B ECI3 |
| | Biological process | Response to chemical | 3 | CRSP THF1 SGT1B |
| | Biological process | Anatomical structure development | 3 | CRSP RPS9M SGT1B |
| | Biological process | Cellular component organization | 3 | WLIM1 THF1 PRMT11 |
| | Molecular function | Transferase activity | 2 | LSMT-L PRMT11 |
| | Molecular function | Hydrolase activity | 2 | PAP1 CRSP |
| $Ca^{2+}$ independent | Biological process | Response to stress | 3 | DGK5 LOX2 PP7 |
| | Biological process | Cellular response to stimulus | 2 | DGK5 PP7 |
| | Biological process | Signaling | 2 | DGK5 PP7 |
| | Biological process | Response to external stimulus | 2 | LOX2 PP7 |

**Supplementary Table S1:** Analysis of GO functional categories for the $H_2O_2$ responsive proteins identified after 10 min of treatment.

| | Pathway database (GO) | GO Functional category | # Proteins | Proteins |
|---|---|---|---|---|
| Strict $Ca^{2+}$ dependent | Biological process | Multicellular organismal process | 6 | ATKDSA2 STR2 CINV1 FIP37 SCE1 FER1 |
| | Biological process | Developmental process | 6 | ATKDSA2 STR2 CINV1 FIP37 SCE1 FER1 |
| | Biological process | Cellular component organization or biogenesis | 6 | WLIM1 ATKDSA2 RPL34A CINV1 XRN3 CSP1 |
| | Biological process | Response to stress | 6 | CINV1 PDF2.2 CPK3 CSP1 FER1 CSD3 |
| | Biological process | Response to chemical | 6 | CINV1 SCE1 CPK3 CSP1 FER1 CSD3 |
| | Biological process | Anatomical structure development | 6 | ATKDSA2 STR2 CINV1 FIP37 SCE1 FER1 |
| | Biological process | Cellular component biogenesis | 5 | WLIM1 ATKDSA2 RPL34A CINV1 XRN3 |
| | Biological process | Reproduction | 4 | ATKDSA2 STR2 SCE1 FER1 |
| | Biological process | Reproductive process | 4 | ATKDSA2 STR2 SCE1 FER1 |
| | Biological process | Developmental process involved in reproduction | 4 | ATKDSA2 STR2 SCE1 FER1 |
| | Biological process | Response to abiotic stimulus | 4 | CPK3 CSP1 FER1 CSD3 |
| | Biological process | Cellular component organization | 4 | WLIM1 ATKDSA2 CINV1 CSP1 |
| | Biological process | Localization | 3 | CPK3 FER1 VPS26A |
| | Molecular function | Oxidoreductase activity | 5 | SCE1 FER1 CSD3 PSBD |
| | Molecular function | Transferase activity | 4 | ATKDSA2 STR2 SCE1 CPK3 |
| | Molecular function | Hydrolase activity | 2 | CINV1 XRN3 |
| | Molecular function | Lyase activity | 2 | PDX12 ASB2 |
| | Molecular function | Small molecule binding | 2 | SCE1 CPK3 |
| Partial $Ca^{2+}$ dependent | Biological process | Cellular component organization | 5 | FREE1 RPL6B CSP3 PME44 GRIP |
| | Biological process | Localization | 4 | FREE1 NUP50A ATPC GRIP |
| | Biological process | Response to stress | 4 | MLP43 CSP3 MES10 PME44 |
| | Biological process | Response to abiotic stimulus | 3 | FD2 CSP3 MES10 |
| | Biological process | Response to external stimulus | 2 | PME44 ATPHOS32 |
| | Biological process | Response to chemical | 2 | MES10 ATPHOS32 |
| | Biological process | Hydrolase activity | 4 | ZFN1 MES10 PME44 ATPHOS32 |
| | Molecular function | Small molecule binding | 3 | 2NADP-ME4 ATPHOS32 |
| | Molecular function | Structural molecule activity | 2 | RPL6B RPL8A |
| $Ca^{2+}$ Indep. | Molecular function | Oxidoreductase activity | 4 | LOX2 CYP97C1 PPOX1 RBCS-1B |

**Supplementary Table S2:** Analysis of GO functional categories for the $H_2O_2$ responsive proteins identified after 30 min of treatment.

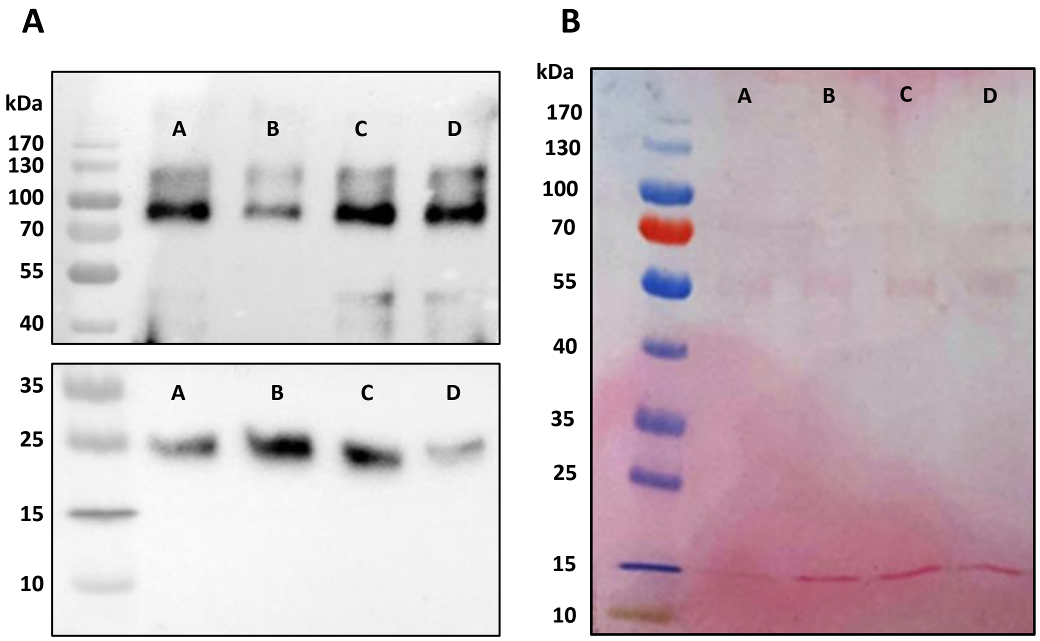


**Supplementary Fig. S2: A:** Comparison of abundance of PAL (top blot) and TIP (bottom blot) in an aliquot of the leaf samples that were used for proteomics analysis. The abundance of PAL and TIP among the different treatments with a duration of 10 minutes (A= Control, B= Stress, C= Inhibitor, D= Inhibitor+Stress) was determined by immunodetection using a specific antibody against PAL and TIP. The Thermo Scientific$^{TM}$ PageRuler prestained$^{TM}$ protein ladder was used to indicate the size of the detected proteins. Due to the distinct sizes of the two proteins of interest the same membrane was used and cut in two before incubation with the primary antibodies. **B**: Ponceau staining of the membrane before incubation with the primary antibodies
